# Optimal competitors: the balance of attraction and choices of mutualists, like pollinators, drives facilitation and may promote crop pollination

**DOI:** 10.1101/2024.10.29.620963

**Authors:** Anna Dornhaus, Alasdair I. Houston

## Abstract

When two species use the same resource, this typically leads to competition, such as when different plants aim to attract the same mutualist pollinators. However, more flowers may also attract more pollinators to an area, such that one or both ‘competitors’ actually benefit from the other’s presence. For example, it has been argued that strips of wildflowers planted next to crops may attract pollinators who ‘spill over’ into the crop. Here we mathematically examine facilitation and competition in consumer attraction. Contrary to previous claims, no accelerating benefits of density *per se* are necessary for facilitation. Instead, under very general assumptions, facilitation can be generated by an imbalance between local competition and joint long-distance attraction of consumers; for example, a low presence of highly attractive ‘wildflowers’ should lead to benefits to a crop. In this mechanism, how pollinator attraction to a patch increases with density of plants is a key factor. Our results generalize to many contexts where local competition may trade off with joint long-distance attraction of consumers, and we show that the exact relationship between competitor density and attraction of consumers can qualitatively shape outcomes, including facilitation or competition.

## Introduction

Our views of ecology and evolution are frequently dominated by competition between individuals and species. Is this justified? It has become clear that in many cases, even organisms that depend on the same resource may facilitate each other’s growth [1], within [2] or across species (in plants [3,4], animals [5,6], or microorganisms [7]). Individuals of other species may prepare habitats or provide additional resources, even while also competing for a joint resource [3,6]. To understand whether one organism competes with or facilitates another overall, we need to understand when these positive effects outweigh the effects of competition. Here we discuss this question in the context of species interactions. We focus onpollination facilitation, a three-species interaction between two plant species and one mutualist (the pollinator), but our model applies to any three-species interaction where one species is a consumer of two other species, whether this interaction is mutualistic or not. We show that, perhaps contrary to intuition, neighbors may be much more frequently facilitators than competitors whenever species depend on attracting consumers. This study thus links the disparate concepts of pollination facilitation, apparent competition (through shared predators), and other types of competition for consumers.

Most flowering plants require animal pollinators to produce seeds, making pollination one of the most well-known mutualist interactions. Most plants are visited by generalist pollinators, who have the potential to visit many species of plants [8–10]. Plants may thus compete for pollinator visits; on the other hand, large patches of flowers may attract more pollinators, potentially generating benefits from such high densities, in other words, facilitation [4,11–18]. While such facilitation has been argued for, empirical data have been inconclusive [6,19–22]. Our model provides some suggestions why this may be the case, i.e. why facilitation may depend on the identity of the species involved and the scale of the study; it will thus allow progress in identifying the causes of context dependence [23].

The case of pollination facilitation in crops has particular applied value [24–26]. Bees provide pollination services of importance to agriculture [27]. However, many bee species are declining [28,29], potentially leading to reduced pollination success [29,30]. This leads to a situation where crop yield may be limited by pollination [25,31]. One proposed solution has been to leave a small section of land next to crops uncultivated (or planted with wildflowers), to attract more pollinating insects, including bees, to the area [25,32]. This may have multiple benefits, including promoting endangered species of both pollinators and wild plant species [33,34] and reducing pests [25]. In particular however, it has been proposed that the presence of diverse plants that share pollinators with a crop may facilitate crop pollination via attraction of additional pollinators – but again with mixed empirical support ([24,25], and possibly [35,36]; no evidence found in [37,38] or a metaanalysis [32]). Some wildflowers are thought to be a natural competitor to crop plants over pollinator visits (sometimes termed the ‘concentrator effect’ because of concentration of bees at the field edge, [14]). If, on the other hand, crop and wildflowers jointly attract a higher total number of pollinators to the area, these might ‘spill over’ into the crop land (also termed the ‘magnet effect’, [11,32]). We use this scenario as the basis of our model, to explore the conditions under which two plants species who share a pollinator may facilitate or compete with each other over pollinator visits.

We derive general insights that apply to any situation in which two species attract the same consumers (whether these consumers are mutualists, as in pollination, or predators). In particular, we demonstrate that the tradeoff between consumer attraction and consumer choice can lead either to competition or facilitation. Our model is inspired by competition over pollinators in crops, but we expand on previous more general theory in plant ecology on pollination facilitation [15,18,39,40]. In previous theoretical work, several authors have demonstrated that pollination facilitation can occur in principle. A primary mechanism examined by previous theoretical studies has been a generalized Allee effect, that is, a situation in which increasing flower density (or nectar availability) from any species leads to an accelerating, i.e. disproportionate, increase in pollinator visits [15,39,40]. In this situation, any additional plants (whether of the same species or not) lead to benefits to an individual seeking to maximize pollinator visits (we include this scenario in Fig. 2f below). We add to these previous models by demonstrating that it is not necessary to assume a general Allee effect (i.e. an accelerating attraction of pollinators with plant density). That is, it is possible for another species to act as a facilitator even when increasing density of *con*specifics has a negative (competitive) effect on individuals. In [15,39], no facilitation was found without a general benefit of density, but our model differs from the above in that we do not assume that across plant species, long-range attractiveness is only a result of the amount of resource provided to the consumer. We also do not make assumptions about population equilibrium and effects of pollinator visits on actual plant reproduction, since, for example in the case of crops, long-term coexistence is not guaranteed (indeed the abundance of both crop and wildflowers is presumably a result of human decisions each year, not plant reproduction). Our results do not depend on how plant reproductive rate depends on pollination, and are thus also applicable to other situations not in equilibrium (e.g. invasive species, [14,30,41,42]). In addition, previous models included other mechanisms that can contribute to facilitation, such as explicit limitations to spatial movement of pollinators [40,43] or flower constancy (limitations to how pollinators switch between flowers, possibly based on cognitive constraints) [22,44]. Even with these additional factors however, facilitation was in previous studies often limited to specific scenarios, e.g. highly abundant competitors [40].

Our study introduces a novel, general, and simple mechanism for facilitation. Specifically, we demonstrate that the balance between attraction of consumers to the area and local choices made by consumers (here, pollinators) critically affects whether facilitation or competition occurs. That is, unlike for example in [15] but like [22,44], we allow the ‘attractiveness’ of plants in terms of increasing the total number of pollinators in a patch to differ from the way in which individual pollinators choose which flowers to visit once in a patch. Empirically, not much is known about how, quantitatively, plant abundance affects consumer attraction and choice; therefore, we examine several different possible functional relationships. Nonetheless, there are many reasons to suspect that pollinator attraction is not simply a linear additive function of the abundance of two species (see also Discussion). The new mechanism for facilitation demonstrated here is likely to apply to other scenarios, including apparent competition, mass flowering, and mimicry, as well as outside ecology to any situation in which two resource providers compete for consumers.

## Model

### Overview of our approach

We formulate a general model for the situation where two resources (which we term ‘crop’ and ‘wildflowers’) compete for consumers (for simplicity called ‘bees’). ‘Crop’ and ‘wildflowers’ can be interpreted to be any two plant species sharing pollinators, such that we focus on the effect of the presence of the second of these two species on the first. We assume that ‘more’ of the wildflowers (*w*) increases the total number of bees in the area (*N*) [16,17,22,39,45–47], but also leads a smaller fraction of these bees to visit the crop (*D*), creating a tradeoff (our notation is summarized in Tab. 1). We term these two processes ‘attraction’ (to the patch) and ‘choice’ (within the patch). We perform both analytical and numerical analyses to explore what determines the amount of wildflowers *w*\* that maximizes the number of bees on the crop, and more generally how the presence of both plant species (i.e. *w* and *c*) affect the number of bees on each. Note that the number of bees on the crop is given by the combination of attraction (*N*) and choice (*D*): *F = ND*.

**Table 1:** Parameters and basic assumptions.

| Parameter name in Model A | Definition | Equivalent in Model B | Explanation |
| --- | --- | --- | --- |
| $w$ | ‘Amount’ of wildflowers | $bp$ | The ‘amount’, or total value of the potential competitor species. This is defined as the quantity in proportion to which pollinators divide up between $c$ and $w$ once in the patch. It should therefore be thought of as including the number of flowers (e.g. density and area covered) and their nectar or resource value (e.g. amount or concentration of nectar). |
| $c$ | ‘Amount’ of crop | $1 - p$ | Analogous quantity to $w$, but for the crop. |
| $N(c, w)$ | Number of bees attracted to local area | $N_p$ | The total number of pollinators in the patch, which we assume depends on the combined ‘amounts’ of wildflowers and crop. We examine different functional forms of $N(c, w)$. |
| $N_c$ | Number of bees attracted by $c$ alone | | The number of bees attracted by $c$ when $w=0$; if $w>0$, this is not necessarily the number of bees visiting $c$, as bees attracted to the area by either plant can end up visiting the other plant, depending on $D$. |
| $D$ | Fraction of bees in local area that visit the crop | $D_p$ | Connects consumer choices to $c$ and $w$ based on an ideal free distribution in the patch: pollinators are dividing up among the plant species in proportion to $c$ and $w$. $D$ can directly be measured empirically. |
| $F$ | Pollinators on crop | $f$ | This is the quantity we want to maximize. In Model A, this is simply the total number of pollinators that visit the crop. In Model B, this is modified by the fraction of area used for crop, such that what is maximized, called ‘yield’ in the text as a shorthand, is proportional both to pollination and area of crop. |
| $w^*$ | Optimal $w$ | $p^*$ | In Model A, this is the value of $w$ that maximizes $F$ for a given $c$. In Model B, we determine the value of $p$ that maximizes $f$. We are interested in determining under what conditions $w^*$ or $p^*>0$, as well as its behaviour under different model assumptions. |
| $A, a$ or $A_c, A_w, a_c, a_w$ | Constants modifying the form of $N(c, w)$ | $A, a$ or $A_c, A_w, a_c, a_w$ | Essentially, the ‘A’ parameters multiply a function that depends on $c$ or $w$ to give the long-range attraction of the respective plants, i.e. how much it attracts pollinators to the patch. The parameters ‘$a$’ modify the shape of $N$ with $c$ or $w$. Both in Models A and in Model B, this parameter affects how quickly the attraction of bees to the area decelerates as $c$ or $w$ increase (implying greater attractivity of rare resources, see Fig. 2d-f and Discussion). Other parameters of $N(c, w)$ explained in text and Supplementary Materials. |

*D* depends on the ‘amount’ *w* of wildflowers and the ‘amount’ *c* of the crop. Throughout, we assume that *D* is simply the fraction of the total resource that comes from the crop (i.e. input matching, [48–51]; Eq. [1]), implying that only the ratio *c*/*w* is important in how bees divide up among resources locally. It is important to note here that the relevant definition of *w* and *c* is ‘the utility of the resources that consumers use to ideal-free-distribute’. In other words, this might be best thought of as a measure of the total quality or quantity of nectar or pollen or whatever resource the consumers are seeking. It is also worth noting that this makes no assumptions about individuals measuring this patch-wide quantity: it is enough that individual consumers are choosing among the resources available those that provide the best return given how other consumers have already distributed themselves [52].

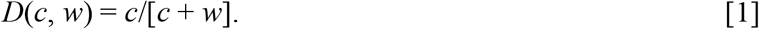

Input matching (and the ideal free distribution) is a general and widely recognized and empirically supported pattern in ecology [51,52]. In prior theory on facilitation, this is also assumed (e.g. in [15], their eq. 7: our *w* represents the same as their *wm*, eq. 6; and in [39]). Studies only diverge from this assumption if they are modeling flower constancy (implying that to some degree individual pollinators choose flowers independently of their resource content, [22,44]).

We specifically model (A) a situation in which the amount of wildflowers is independent of the (fixed) amount of crop *c*, and (B) one in which the total area available is fixed, and only a fraction of the area is devoted to wildflowers (with the rest taken up by the crop). This resembles a situation where the two plant types compete for a common resource other than the pollinators (as in [15,39]).

The complete code for the numerical calculations and to produce the figures was written in R (version 4.4.2) using RStudio and the packages ‘scales’ [53] and ‘viridis’ [54], and is available on github (https://github.com/dornhaus/spillover).

## Results

### Model A: Independent *w*

We first make a very general argument about the optimal amount of wildflowers, i.e. the *w*\* at which *F* is maximal for a given *c*. We make no specific assumptions about the shape of *N* other than that it is decelerating (rate of pollinator attraction eventually slows, [16,18,46]) with *w. F* is maximal at

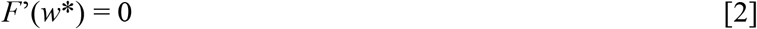

So

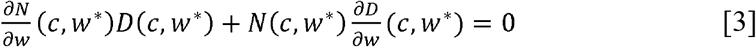

Given eq. [1], it follows that

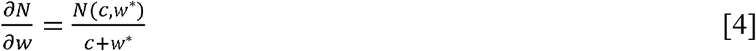

Similarly to the graphical solution for the marginal value theorem [55], we show graphically (Fig. 1a) that this implies that *w*\* increases as *c* increases, as follows. We plot an x-axis with *w* (the ‘amount’ of wildflowers) increasing from the origin to the right, and *c* (the ‘amount’ of crop) increasing from the origin to the left (Fig. 1a). We plot *N*(*w*), that is, the total number of bees attracted to the patch as a function of *w*; we assume that this is an increasing, but decelerating function (we use eq. [5] to generate the plot but the precise shape is irrelevant as long as it is continuous and eventually decelerates). We then draw a tangent, i.e. a straight line starting from -*c* on the x-axis to the point where it touches the function *N*(*w*). At this tangent point, the slope of *N, N*’(*w*), is equal to the slope of the tangent, *N*(*c, w*)/(*c*+*w*). Since this satisfies eq. [4], and thus maximizes *F*, the value of *w* at this point is *w*\*, the optimal amount of wildflowers to maximize the number of bees on the crop (second vertical dashed line in Fig. 1a). Below we discuss some limited conditions (numerical parameter values) under which *w*\* may not be >0.

**Fig. 1:**
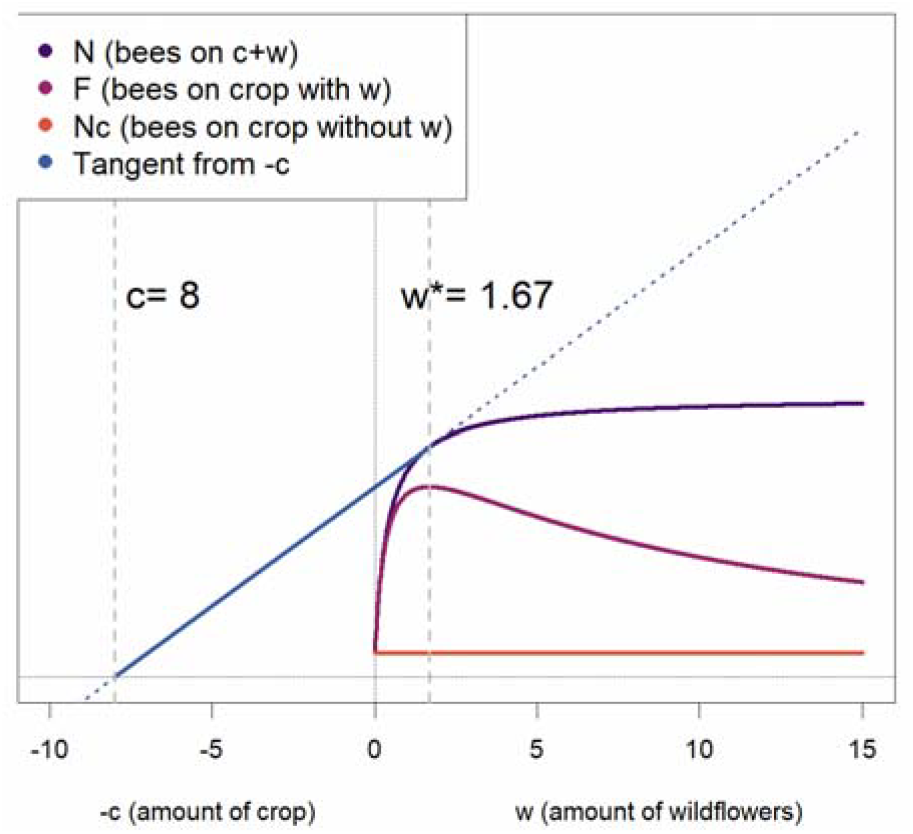
Graphical solution to finding *w\** (compare to marginal value theorem, [55], and see text for full explanation). The number of bees on the crop (*F*) peaks at a low but non-zero presence of the competitor (*w*), even when the total number of bees attracted to the patch (*N*) is never accelerating with *c*+*w*. This argument does not depend on the shape of *N*(*c*,*w*), as long as it decelerates in *w* and *N*(*c*,0) (which we call *N*_c_) is not too large. The quantitative size of the facilitative effect of *w* depends on specific assumptions (see Supplementary Materials; for the purpose of this plot, we assumed *N* is given by eq. [5], with parameters *a*=0.4, and *A*=10).

**Fig. 2:**
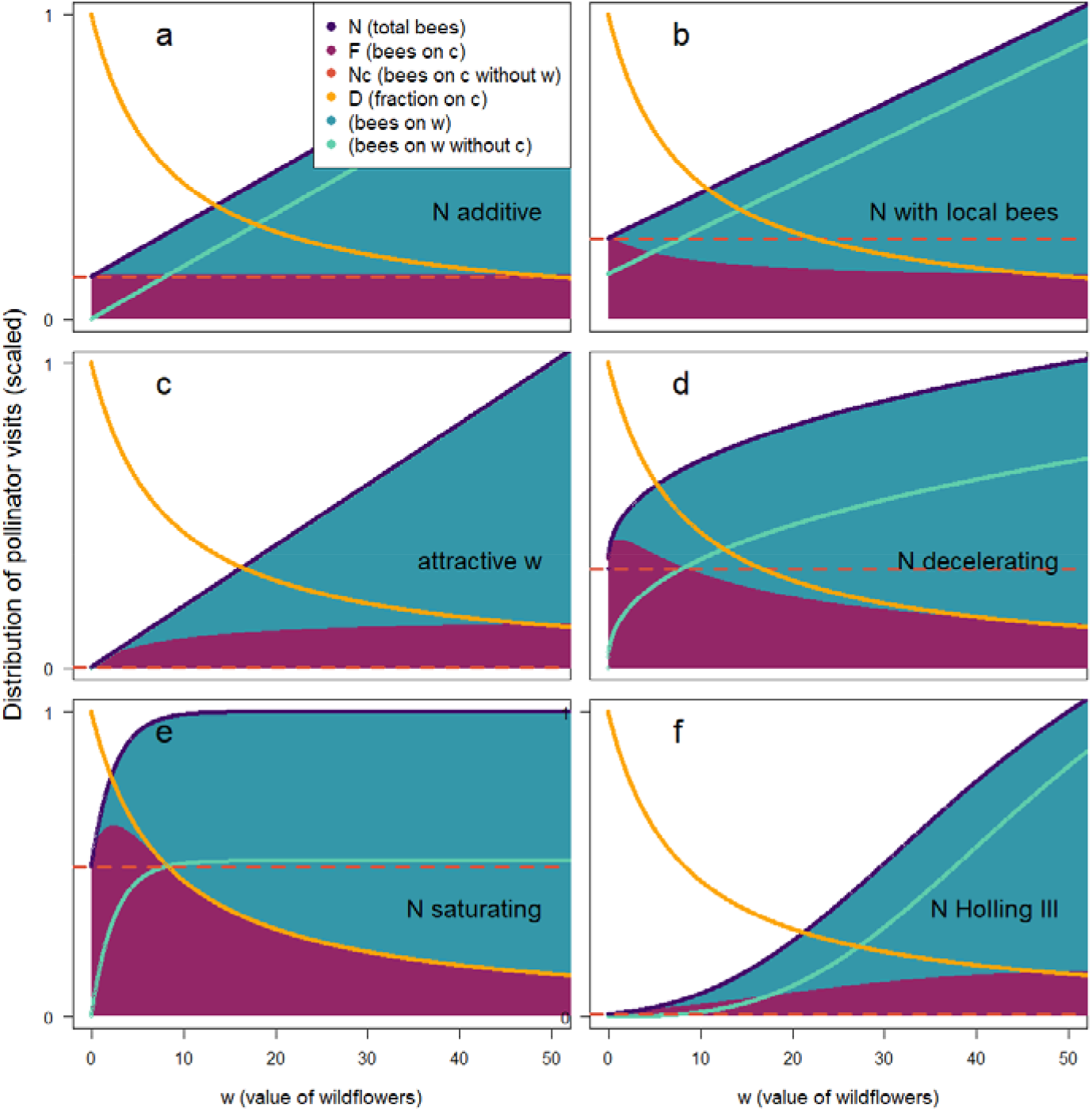
The qualitative effect of different relationships for how the total number of bees attracted to a patch, *N*, depends on *w. D*, the fraction of pollinators on the crop, behaves the same in all panels as it only depends on *c* and *w* (not *N*). Wherever *F* (bees on crop, *F*=*D*\**N*) is above *N*_*c*_ (bees on crop without *w*), facilitation is occurring (*c* is held constant in all panels). *w* has no effect on *F* in (a), because each plant attracts exactly the number of pollinators that choose it (*N* is simply a linear function of *c* and *w*). In (b) however, *w* competes with the crop (reduces *F*), because the plants compete for the ‘local bees’. In (c), a highly attractive *w* (i.e. a plant which attracts bees disproportionally to the patch given how bees in the patch choose it) facilitates the crop (increases *F*). In (d,e,f), facilitation may or may not occur: in all these cases, the attraction of bees to the patch does not behave as the choice of bees in the patch. In all panels, *N* is scaled to 1 at *w*=50 (and *F* and the analogous number of bees on the wildflowers are scaled accordingly), to be able to qualitatively show all relationships. Parameters are set to *a*=0.4, *A*=10, *L*=10, *r*=0.02, *y*=3, *g*=1.

#### The effect of attraction: different forms of N

Our graphical solution in Fig. 1 is based on a situation where *N* is decelerating with *w*. That is, the number of bees attracted increases more and more slowly with an increasing amount of wildflowers (empirically demonstrated for pollinators or herbivores in [16,18,46,56]). However, the outcome for either species depends critically on how exactly *N* behaves. If the number of bees is simply proportional to the sum of crop and wildflowers available (*N*=*g*(*c*+*w*), where *g* is a positive constant, Fig. 2a), then there is neither competition nor facilitation: each plant attracts exactly the number of bees that visit it (*F* stays constant over *w*). If, in addition to the above, there is a constant number *L* of local bees in the patch regardless of *c* and *w*, and the additional number of bees attracted to the area remains *N*=*g*(*c*+*w)*, then competition for these local bees occurs, and the number of bees visiting the crop is reduced by any presence of wildflowers (Fig. 2b).

However, if wildflowers are disproportionately good at attracting pollinators (*N*=*Agw*+*gc*, Fig. 2c), or if either crop or wildflowers show diminishing returns in attracting pollinators (whether that is merely decelerating, as in Fig. 2d, or saturating, as in Fig. 2e; we examine these cases analytically in more detail in the Supplementary Materials, as models A.1, A.2, and A.3), then the number of bees attracted by wildflowers does not behave as the fraction of bees that then choose wildflowers once they arrive in the area. As a result, the number of bees visiting the crop (*F*) does not stay constant with the amount of wildflowers present, and facilitation or competition may result. For the case with diminishing returns (Figs. 2d,e), the intuitive interpretation is that with diminishing attractiveness of a flower type with the amount of that flower present, the maximal number of pollinators attracted per flower is at low abundance, and thus the maximal total number of pollinators in a patch is achieved by a mix of both flower types. Past models have emphasized situations in which there is initially an acceleration of *N* with *c*+*w* [15,39], such as in a sigmoid relationship (cf. Holling type III functional response, *N*= (*r*(*c*+*w*))^y^/(1+((*r*(*c*+*w*))^y^), Fig. 2f). In such a case, facilitation is not surprising, and results from increasing density of either conspecifics or heterospecifics: any added plants attract disproportionately more bees (Fig. 2f). However, as we show here, such acceleration is not a necessary condition for facilitation (cf. Fig. 2c-e).

Overall, these calculations demonstrate that the facilitative effect in our model comes from the imbalance between the attraction of bees to the local patch generated by the combination of the two flower species, and the choice those bees make once they have arrived at the patch. Eq. [1] assumes that there is some way in which the ‘amount’ of each flower species (here called ‘*c*’ and ‘*w*’) can be quantified such that bees in the patch divide themselves among the flowers in proportion to this ‘amount’ (whether this is number of flowers, quality or quantity of nectar, or some other measure). Fig. 2 illustrates that how these ‘amounts’ relate to attraction of bees (pollinators) to the patch drives whether facilitation or competition for pollinators occurs.

#### Model A.1: N as decelerating function of w

To better assess the quantitative expectations for *w*\*, we assume a particular shape of *N*:

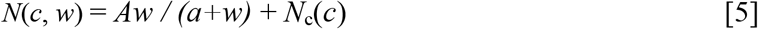

This gives a relationship between *N* and *w* that is monotonically increasing but with diminishing returns (slope) as *w* increases (as Fig. 1, and in Fig. 2d), and thus is a specific case of the ‘Model A’ presented above. Note that we analytically demonstrate that the outcome is qualitatively identical for two alternative functional forms of *N*(*c*,*w*) in the Supplementary Materials, including a power function (Model A.2, identical to [15]) and an exponential, saturating function (Model A.3).

At the *w\** that maximizes the number of bees on the crop (i.e. *F*), the partial derivative of *N* should be equal to

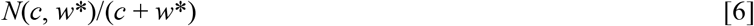

(cf. Fig. 1f). The partial derivative of eq. [5] together with eq. [6] gives as the optimality condition

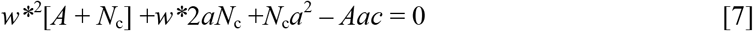

This quadratic equation can be rearranged to give

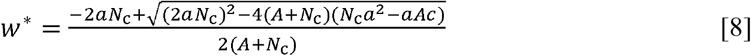

We examine the numerical consequences of the parameters *a* and *A* on this result in Supplementary Materials. Generally, facilitation occurs under almost all circumstances, for both species. For the *N*(*c*,*w*) in eqn [5], the only requirement is that *c* > *c*_0_, where *c*_0_ is the value of *c* at which *w*\* reaches zero. Putting *w*\* = 0 in eqn [7] gives *c*_0_ = *N*_c_*a*/*A* (note that there is always a positive *w*\* in model A.2, see Supplementary Materials). The highest benefit for the crop (i.e. the highest *F*) results if the second species (the wildflowers) are more attractive to pollinators from a distance (high *A*) for the same ‘value’ locally (i.e. per *w*) (Fig. S2d-f).

### Model B: Fixed total area, such that *c* and *w* trade off

Alternatively, we might assume that the total area in which either crop or wildflowers can be planted is fixed. Thus, increasing the amount of wildflowers requires a corresponding reduction in the amount of crop. We designate *p* as the proportion of total area devoted to wildflowers; the proportion of area devoted to crop is thus 1 - *p*. We further designate *b* as the relative value (to bees) of wildflowers per area compared to the value of crop per area, such that the choice of bees once in the patch, *D*, remains as in eq. [1]. If we standardize the ‘value’ of the crop at *p* = 0 to 1, then

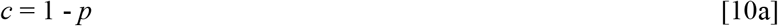

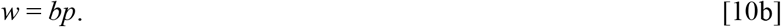

The number of bees and the fraction on the crop are now functions of *p* so we change our notation to *N*_*p*_*(p)* and *D*_*p*_*(p)* to reflect this. We expect *b* > 1 as we assume that wildflowers are more attractive to bees than the crop is (per area). As a result, we expect that increasing *p* will increase *N*_*p*_, decrease *D*_*p*_, and decrease the area available for the crop. We want to maximize ‘yield’ (*f*), which we take to depend linearly both on the number of bees visiting the crop (*N*_*p*_*D*_*p*_) and the crop area (total area multiplied by 1 - *p*). While we assume that *f* needs to be maximized to benefit the crop, calling this quantity ‘yield’ is only a shorthand, expressing the assumption of pollination limitation and of linear dependence on area; we do not model fruit production or other processes affecting actual yield. Thus,

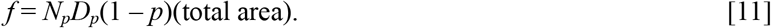

At maximal *f, f*’ = 0. Note that *N*_*p*_ and *D*_*p*_ both depend on *p*, but ‘total area’ is a scaling factor that does not affect the optimum. The differential (with respect to *p*) of the product of the three terms that depend on *p* is

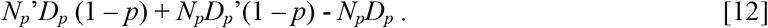

This equals 0 at *p* = *p*\*, giving

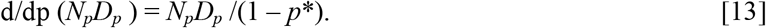

Analogously to Model A, we assume a form for *N* that implies a decelerating effect of resource availability on the number of bees attracted:

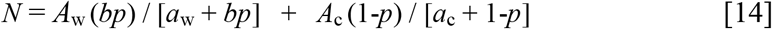

We numerically find *f* for different parameter values, as well as the resulting *p*\* (Fig. 3; see Supplementary Information for numerical calculation of the effects of all parameters, including the case where the parameters *A*_w_ and *A*_c_, and *a*_w_ and *a*_c_, respectively, are not the same). As in Model A, the consequence of decelerating attractiveness of each species with density is that the maximal number of pollinators gets attracted to the patch at some mixture of the two species, and this leads to a benefit for each species (in terms of pollinator visits) from at least a small amount of the ‘competitor’. The largest optimal ‘competitor’ presence is achieved with highly attractive (high *A*_w_), high quality (high *b*) ‘competitor’ species, whose own attractiveness is not steeply decelerating (lower *a*_w_) (Fig. S3).

**Fig. 3:**
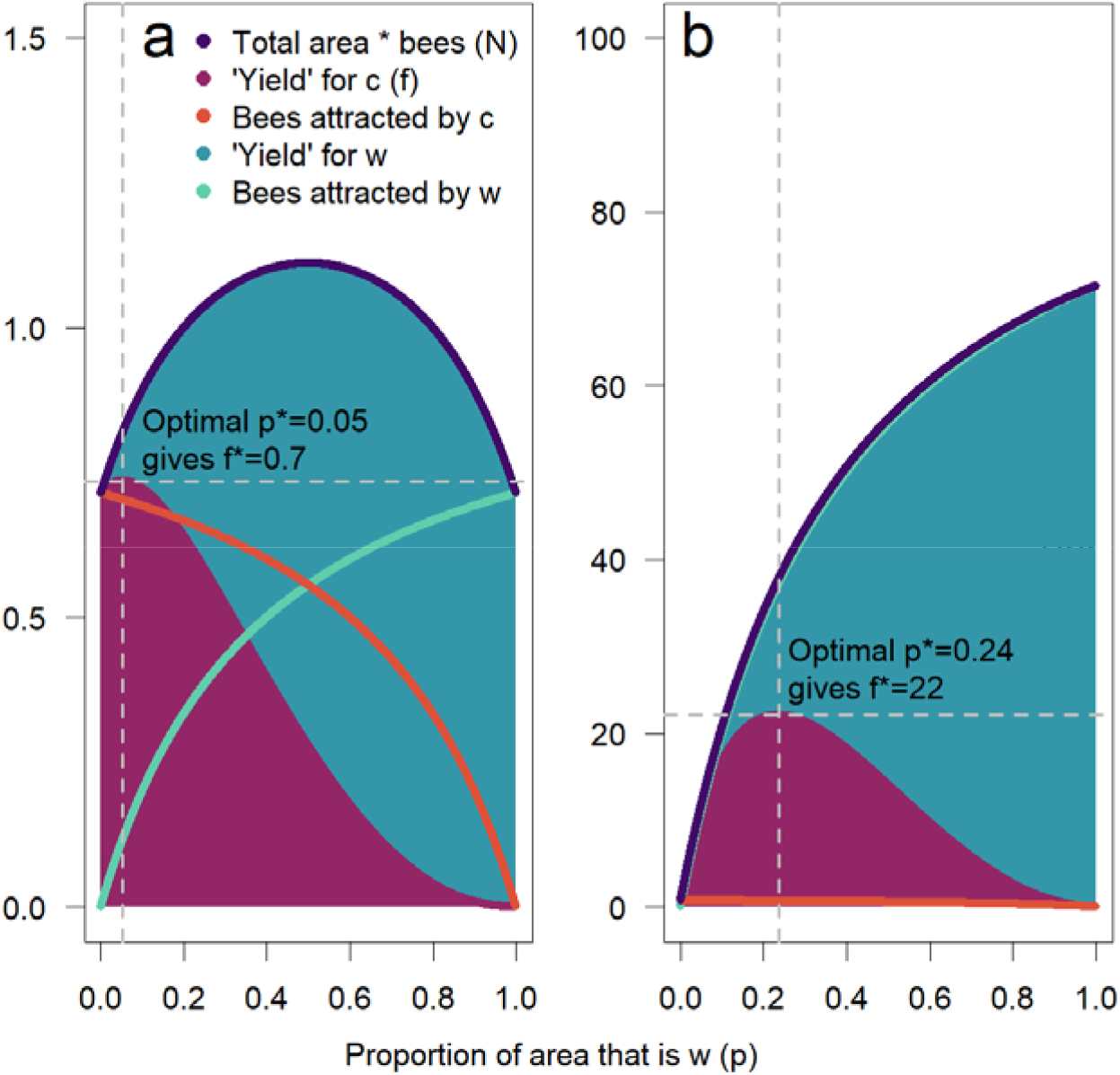
The ‘yield’ (area multiplied by number of bees) for each plant in Model B (total area fixed). (a) Even assuming both plants are exactly equal in attractiveness and value per area (*A*_w_=*A*_c_=1, *b*=1, *a*_w_=*a*_c_=0.4), because of the decelerating returns in attraction from each resource individually (eq. [14]), the total number of bees is highest at *p*=0.5, and ‘yield’ for the crop is highest at a *p*\*>0. Note that the *A* and *a* parameters relate to how each species attracts pollinators to the patch, whereas *b* relates to the relative ‘value’ per area of each plant (thus affecting choice of bees once in the patch). Therefore, because bees choose both resources in proportion to their value, but attraction to the patch by each species decelerates with its value (or abundance), each plant achieves a net benefit from a small presence of the other plant species (here, *f* peaks at *p*=0.05). (b) If *w* is much more attractive than *c* (*A*_w_ =10, *A*_c_=1), the total number of bees attracted is almost exclusively driven by *w* (and thus increases with *p*). But because *f* is determined by 1-*p* as well as the number of bees on *c*, there is a tradeoff: more *w* means more pollinators but less area for the crop. Nonetheless, a more attractive *w* ultimately leads to a benefit for *c* (*f*\* is higher for higher *A*_*w*_; note difference in y-axis scale between panels).

## Discussion

What makes a neighbor a competitor? When success depends on the attraction of consumers (such as mutualist pollinators), the presence of other individuals that contribute to such attraction may lead to benefits, whether they are conspecifics or heterospecifics. Previous theory had shown that when pollinator attraction accelerates with density of plants, higher density is beneficial to individuals [15,39]; this is also termed the ‘Allee effect’, particularly in the animal literature [6], and is a specific example of facilitation [3]. Here we show that such disproportionate benefits of density (i.e. accelerating attraction of pollinators) are not necessary for one species to facilitate pollination (or visitation by any consumers) of another. Instead, we show generally that the balance of competition and facilitation in pollination can be driven by the balance (or imbalance) between attraction-to-the-area and the choices-made-locally of consumers. Such an imbalance can arise in a number of ways (see below), but seems particularly likely when consumers seek variety, i.e. are attracted especially by more ‘rare’ resources. In this and many other cases, as the abundance of the potential competitor plant increases, attraction of additional pollinators to the area and number of pollinator visits to this ‘competitor’ may both increase, but need not behave in the same way (see also [22]). Whenever the marginal increase of consumers attracted by an additional heterospecific plant is higher than the marginal decrease in consumers choosing the focal plant species, facilitation occurs. As we show, the commonly-assumed (and empirically demonstrated, [18,46,56]) decelerating marginal attractiveness of additional plants of the same species therefore is likely to enable facilitation to occur. This is only limited if *N*_c_ is large and *c* is small, i.e. if the focal species already attracts many consumers to the patch but is poor at providing value to them once they are there.

### The difference between attracting pollinators to the area and local choices of pollinators

Why might we expect that attraction to the area is not the same as how consumers choose locally? There are four types of processes that may play out differently across these spatial scales. First, how flowers are perceived from a distance is determined by different aspects than how flowers are perceived when near (e.g. olfactory vs visual signals, [57,58]), thus potentially leading to differences in attractiveness between plant species across spatial scales. Second, pollinators may often require and seek out micronutrients, which can attract them over some distance, even when the proportion of visits that bees need to make to flowers offering those nutrients is not large [59–61]. Third, ‘attraction’ may occur over different (longer) timescales than local choices, for example when bees are attracted to nest near particular species [26,62] and then forage primarily locally, or if bees make long foraging trips only to some patches but once there visit almost all available flowers [40,63,64]. In one system, extremely fast local reproduction of the pollinators combined with slow (if any) movement between patches may have led to extreme pollination facilitation (and concomitant evolution of mass flowering, [43]). While we don’t explicitly model such temporal dynamics, they can lead to different effects of the abundance of a particular plant species on attraction of bees vs. choices by bees in the local area. Fourth, we frame the ‘local area’ that bees are attracted to as a spatial location, but it may be viewed as any set of flowers that bees, once attracted, are faithful to: for example, a learned color (‘flower constancy’, [65,66]). In this case ‘attraction’ is about how likely bees are to become flower-constant on the particular color or flower type, while ‘choice’ reflects (as before) how bees that are already committed (flower constant) on this type choose among the different species [44,66]. In all these cases, the number of pollinators available in a patch (*N* in our model) behaves differently with changes in abundance than the proportion of bees visiting a focal plant species (*D* in our model), leading the ‘neighboring’ species to become either a competitor or facilitator. Moreover, a non-linear attraction of bees can lead to effects where both plant species may facilitate (or harm) each other (see Fig. 2). In our focal model, we assume a decelerating attractiveness (of each plant species), which is what one would expect for example if plants vary somewhat in the type of resources provided (e.g. pollen vs. nectar, amino acids, or other micronutrients), and bees seek balanced diets. In such cases, bees experience decreasing marginal returns of each resource type, an extremely common situation in foraging, resulting in facilitation.

### When is facilitation likely in practice?

Our model leads to several specific conclusions. First, plant species that lead to an increase of pollinators in the patch are more likely to facilitate others. These might be species offering particular micronutrients, species with prominent long-distance signals, species that encourage pollinators to travel long distances to a patch, or species promoting pollinator reproduction. Any attraction of pollinators to the patch increases the total ‘amount of pollination’ received by the plants in the patch, and thus prevents competition from being inevitable, as well as creating the possibility of mutual facilitation. Second, our model shows that subtle variations in the functional relationship of attraction and choice with plant abundance have qualitative effects on species interactions (and it is thus critical to identify which of the regimes in Fig. 2 applies). Frustratingly for the empiricist, relationships that look quantitatively similar may have qualitative consequences [68–70]. Several studies have shown support for an increasing, decelerating attraction of pollinators to a patch (*N*), albeit with the possibility of an accelerating *N* at very small patch sizes [16–18,22,24,56,67]. It is thus not unlikely that plants experience both intra- and interspecific facilitation at such very small patch sizes (Fig. 2f). At a larger patch size however, the rarer species is likely to attract disproportionately more pollinators (because most resources yield diminishing returns for consumers). In such cases, the more common species will benefit from the presence of the rarer one. This is contrary to the common notion that it is only the rare species that benefits [16,22]. However, that notion probably derives from the common empirical scenarios of extremely small total patch sizes. Nonetheless, and third, our model also shows that an intermediate abundance of both species can lead to maximal visits of pollinators per plant for both species, and in fact this is a likely outcome. The quantitative amount of facilitation generated, however, is highest for the species that is abundant, yet in relation to its abundance not particularly good at attracting pollinators from a distance.

### ‘Apparent’ facilitation

The model presented here provides a general framework for thinking about species interactions other than pollination mutualism at different scales simultaneously. Most obviously it may apply to other mutualistic interactions, for example beneficial microbes and fungi [71]. However, it also applies to antagonistic interactions. If we imagine the ‘bees’ in our model as predators, the conclusions from our model hold: the bees play the role of ‘shared predator’ (whether they consume the flowers or just their nectar and pollen is, after all, immaterial to the bees’ choices), and the different flower species are the ‘prey’. It follows that the presence of a second prey species in a patch may lead to either increased predation on a focal species (the equivalent of pollination facilitation) or decreased predation (the equivalent of pollination competition, which in the case of predation would be a facilitative effect) [67,72]. Whether predation pressure is increased or decreased by additional prey species will thus depend, analogously, on the ability of these other species to attract predators to the area (or to the type of prey, analogous to flower constancy, discussed above) relative to how likely predators are to choose that prey once they encounter it [73].

In ecology, the term ‘apparent competition’ refers to the phenomenon that two (prey) species may affect each other’s population dynamics by virtue of their effects on a shared predator [74]. Typically, models of apparent competition focus on the effects of the prey on the actual population growth of the predator (and in turn how this affects the growth of prey populations), rather than the sub-population-level processes modeled here (of patch and item choice of predators). Our model does not consider population growth for either crop, wildflowers, or bees. Nonetheless, the process described by our model can also explain an effect of one prey species on another via a shared predator: in cases where any two prey species share a predator that may be attracted to a local area, a process like ‘apparent competition’ or ‘apparent facilitation’ could occur. As per our model, this will depend on the abundance of both prey types as well as how it relates to their ability to attract predators to the area. We argue here that ‘spillover’ or pollination facilitation is quite general, and indeed this has been analogously assumed in the apparent competition literature (i.e. increased predation as a result of similar species, [73]). However, just as plants may occasionally compete for pollinators (see Fig. 2b and outcomes at high *w*), predators may result in ‘apparent facilitation’, i.e. one species of prey may draw predators away from another, leading these two prey species to ‘apparently’ facilitate each other (rather than displaying ‘apparent competition’).

For the particular case of crops and wildflowers, it should be noted that our model also applies to herbivores, i.e. potential pests. That is, if wild plants share herbivore consumers with the crop, they may attract such herbivores, which then ‘spill over’ into the crop land. Data is needed to determine the degree of such spillover; the attractiveness of any particular plant to herbivores is not likely to be the same, or even scale in the same way with abundance, as the attractiveness to pollinators. What also bears mentioning is that previous data show spillover at yet another trophic level: wildflower margins next to crops may lead to higher abundance of insect predators, i.e. pest control [25].

### Conclusion

Our model derives an important and general principle that has not previously been recognized in the study of pollination facilitation. Broadly, it is that when it comes to consumers as a resource, the presence of other individuals, both conspecific and heterospecific, affects the abundance of that resource both in positive (long-distance attraction) and negative (local competition) ways. When these effects cancel each other out, a neighbor is neither competitor nor facilitator, as each individual attracts exactly the amount of resource ‘consumed’ (Fig. 2a). Under a broad set of ecological circumstances however, we expect the marginal attraction of consumers to be higher for the rarer resource (what we model as decelerating *N*, Fig. 2d,e). We show that such circumstances are likely to lead to facilitation.

In plant ecology, pollination facilitation has been much discussed (e.g. in models [15,39,40,75] or empirically [11,18–21]). In contrast to previous models however, our study demonstrates that facilitation can occur even under very minimal assumptions, namely, just two ecological principles: the numerical response [76,77] and an ideal free distribution [21,51,52], both standard approaches when describing how consumers select between resources. Facilitation or competition can thus occur without the need to assume more complex processes like resource competition [15,39], flower constancy [22,40,44], nonlinear relationship of pollination and reproduction [78], or specific spatial or temporal scales [32,34,40]; we also do not require that pollinator attraction is an accelerating function of plant abundance [15,39]. Such other processes may of course affect the balance of competition and facilitation in terms of plant reproduction. Nonetheless, when it comes just to attention from consumers, our model demonstrates that facilitation does not depend on such very specific assumptions, and many of the above processes can be seen as special cases in our model (specific parameter constellations, Fig. S2).

The key novel insight from our study is that an imbalance between attraction to the local area and choice of pollinators within the area alone can generate pollination facilitation between plants, or generally between species competing for consumers. Ultimately this tension between attraction at the larger spatial scale (e.g. the field, the patch, or a set of preferred resources/prey) and competition at the smaller scale (e.g. individual plants or prey, wildflowers vs a crop plant) affects the way in which different resources affect each other’s consumer visits, with implications not only in pollination but species interactions across ecology (predator-prey interactions, mimicry) as well as other fields interested in the relationship between resources and consumers.

## Acknowledgements

The authors thank Judie Bronstein and John McNamara for comments on earlier versions of this manuscript.

## Supplementary Materials

The supplementary materials contain the following sections:

1. Models A.1-A.3: specific formulations of *N* decelerating with *c* and *w*
  a. Illustrating the effects of different amount of each species on facilitation
  b. Illustrating the effects of different parameter values
2. Model B: effects of parameters on attraction and choice for fixed area

### 1. Models A.1-A.3: specific formulations of N decelerating with c and w

#### a. Illustrating the effects of different amount of each species on facilitation

Throughout our paper, we assume that local choices of consumers are given by *D*(*c, w*) = *c*/[*c* + *w*] (eq. [1]). Another way of saying this is that *D* describes the proportion of pollinators on the plant species *c*, out of the total who are present in the patch. This can be directly empirically measured.

We explored the effect of widely differing relationships between *N* and *c* and *w* in Fig. 2, demonstrating the general phenomenon that if the total number of pollinators attracted (*N*) does not behave as *D* does with changing ‘amounts’ of each resource, either competition or facilitation may occur. Our analytical result, however, showed generally that facilitation can occur for any decelerating function of *N* with *w*.

Here, we formulate three different specific models that show this qualitative behaviour (increasing but decelerating *N* with *w*), and then numerically explore how the quantitative outcome (e.g. the amount of facilitation and the optimal ratio of *w* and *c*) depend on specific assumptions about parameters in these models (note that although we use the same parameter names, these have different effects in each model and not the same biological meaning; see next section).

Model A.1:

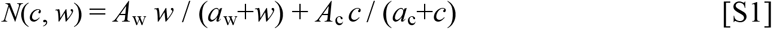

Model A.2:

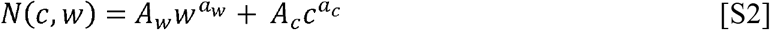

Model A.3:

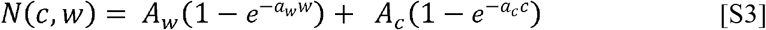

All three models show monotonically increasing, but decelerating *N* with either *w* or *c* (Fig. S1). When *N* is decelerating, this implies the number of bees attracted to the patch per plant is higher at lower abundance (*w* or *c*), and thus the total number attracted by both flower types is highest at intermediate *D*. As Fig. S2 shows, in models A.1 and A.3 the number of pollinators attracted (*N*) saturates both as *w* increases (and *D* decreases) and as *c* increases (shown in top row of Fig. S2). As a result, *D*(*w\**) increases with *c* in A.1 and A.3. In A.2, interestingly, *D*(*w\**) stays constant with *c* (although this requires an increasing *w\** to match the increasing *c*, and requires *a*_w_=*a*_c_). However, regardless of the value of *c*, facilitation occurs at intermediate *D*.

In all three models, regardless of specific parameter assumptions, facilitation can occur (*w*\* > 0) at least if *N*_c_ is small and *c* is large, in other words, when the ‘crop’ is not very effective at attracting pollinators from a distance. Mathematically, if *N*_c_ = *N*(*c*, 0) = 0 (in all models), and always in model A.2, we have the usual Marginal Value Theorem case: there is always a *w*\* > 0 if *c* > 0. In Models A.1 and A.3 there can be a *c*_0_ > 0 such that *w*\* = 0 if *c* < *c*_0_. In A.1 there is an explicit equation for c_0_:

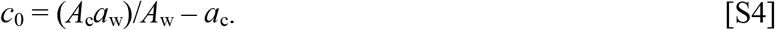

this equation can be obtained from the equation *c*_0_ = *N*_c_*a*/*A* together with *N*_c_ = *A*_c_*c*/(*a*_c_ + *c*).

In Model A.3, *c*_0_ satisfies

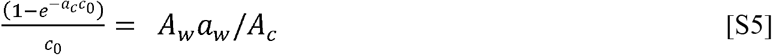

Thus, in both Model A.1 and A.3, a higher ratio of *A*_w_/*A*_c_ makes facilitation more likely, but facilitation can occur even if this ratio is <1, as long as *c* is higher than *c*_0_. In other words, even ‘less attractive’ species may still facilitate pollination of a focal species, as long as there is a large amount of the focal species present. The amount of pollinators gained, however, is only likely to be significant if the wildflowers are more effective than the focal species at attracting pollinators (*A*_w_>*A*_c_).

**Fig. S1:**
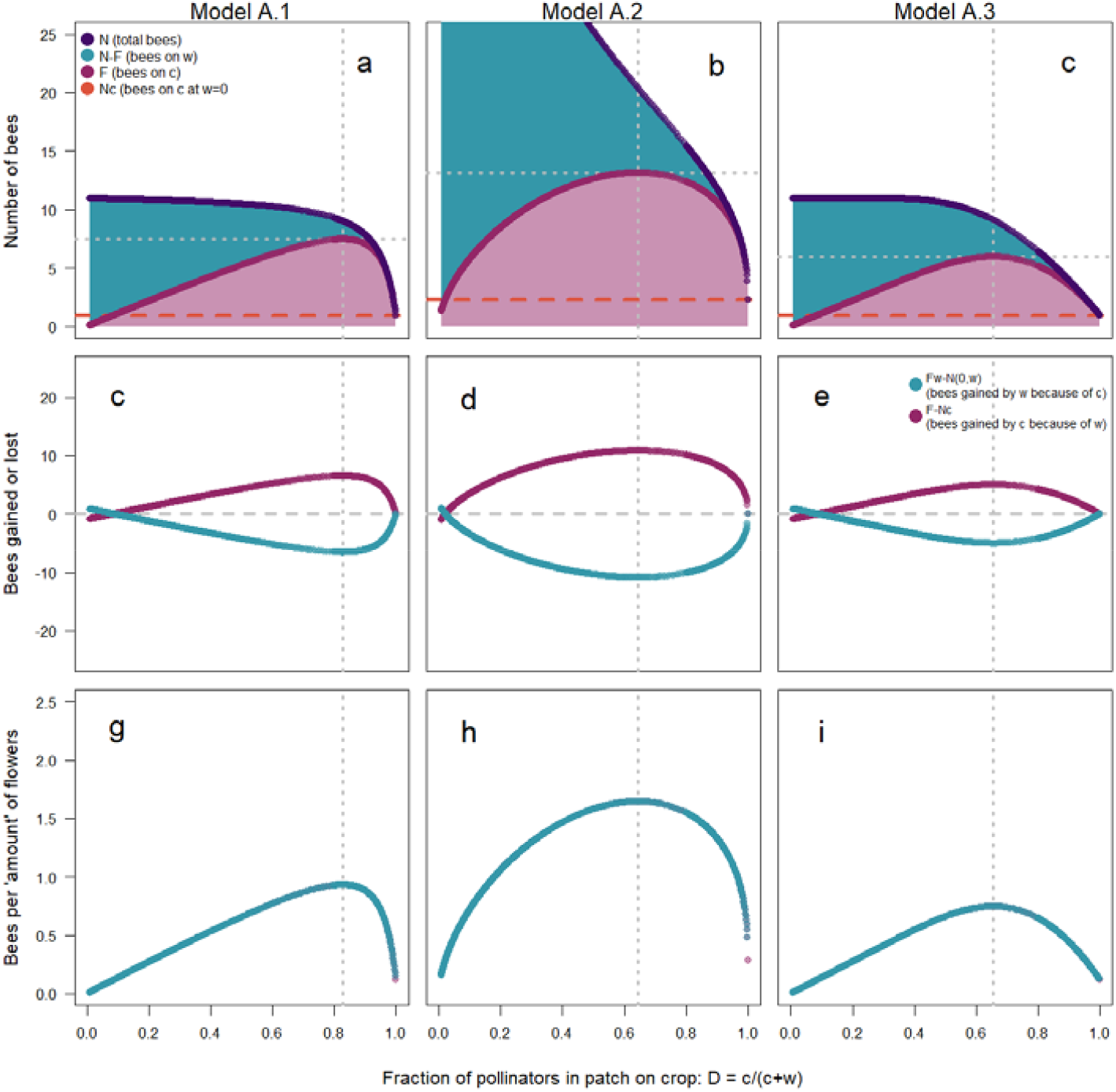
Three specific ways of modeling attraction of pollinators to the patch with changing *w* (a, c, g: model A.1; b, d, h: model A.2; c, e, i: Model A.3). Here, we show *D* on the x-axis, which is the empirically measurable fraction of pollinators in the patch that visit *c* instead of *w*. As *w* increases, *D* decreases. Panels a, b, c show the total number of bees attracted to the patch (blue) and the number visiting the crop (red), similar to Figs. 1 and 2 (except as a function of *D* not *w*). Panels c, d, e show how many bees are gained and lost respectively by the crop and the wildflowers as a result of the presence of the other species, in total. However, by definition the number of bees on crop and wildflowers per ‘amount’ of these flowers (i.e. per *c* and *w*, respectively) is the same (shown in panels g, h, i). This implies that at *w\** (vertical stippled lines), both species can expect the maximal number of bees per individual flower – and since *D*(*w\**) is always <1 (and *w\**>0), facilitation for both species occurs. Parameter values: *c*=8, *A*_w_=10, *A*_c_=1, *a*_w_=*a*_c_=0.4.

#### b. Illustrating the effects of different parameter values

In all models, *A*_*w*_ and *A*_*c*_ are parameters reflecting the (long-distance) attraction of pollinators to the patch relative to the choices made by pollinators in the patch (*w* and *c* respectively). Mathematically they are factors multiplied by a function of *w* or *c* to give *N*. As a consequence, higher *A*_*w*_ strongly increases *F* and thus the benefits gained by *c* from the presence of *w*; in Fig. S2, panels d-f also show that higher *A*_*w*_ decreases the optimal *D*(*w*\*) (‘optimal’ from the point of view of maximizing *F*, i.e. the benefit to the crop in terms of pollinator visits). The parameters *a*_w_ and *a*_c_ on the other hand vary in their role across the three models, although they always affect the shape (curvature) of *N*(*c*,*w*). The shape of *N*(*c*,*w*) as reflected by the value of *a*_w_ has widely varying effects on both *F* and *w\**, demonstrating that the details of the functional form of *N*(*c*,*w*) can have strong effects on outcomes (Fig. S2, panels g-I; for a discussion of this general point see also [68]). The parameter *a*_c_, however, has little effect while *c* is held constant, as in panels j-l in Fig. S2.

**Fig. S2:**
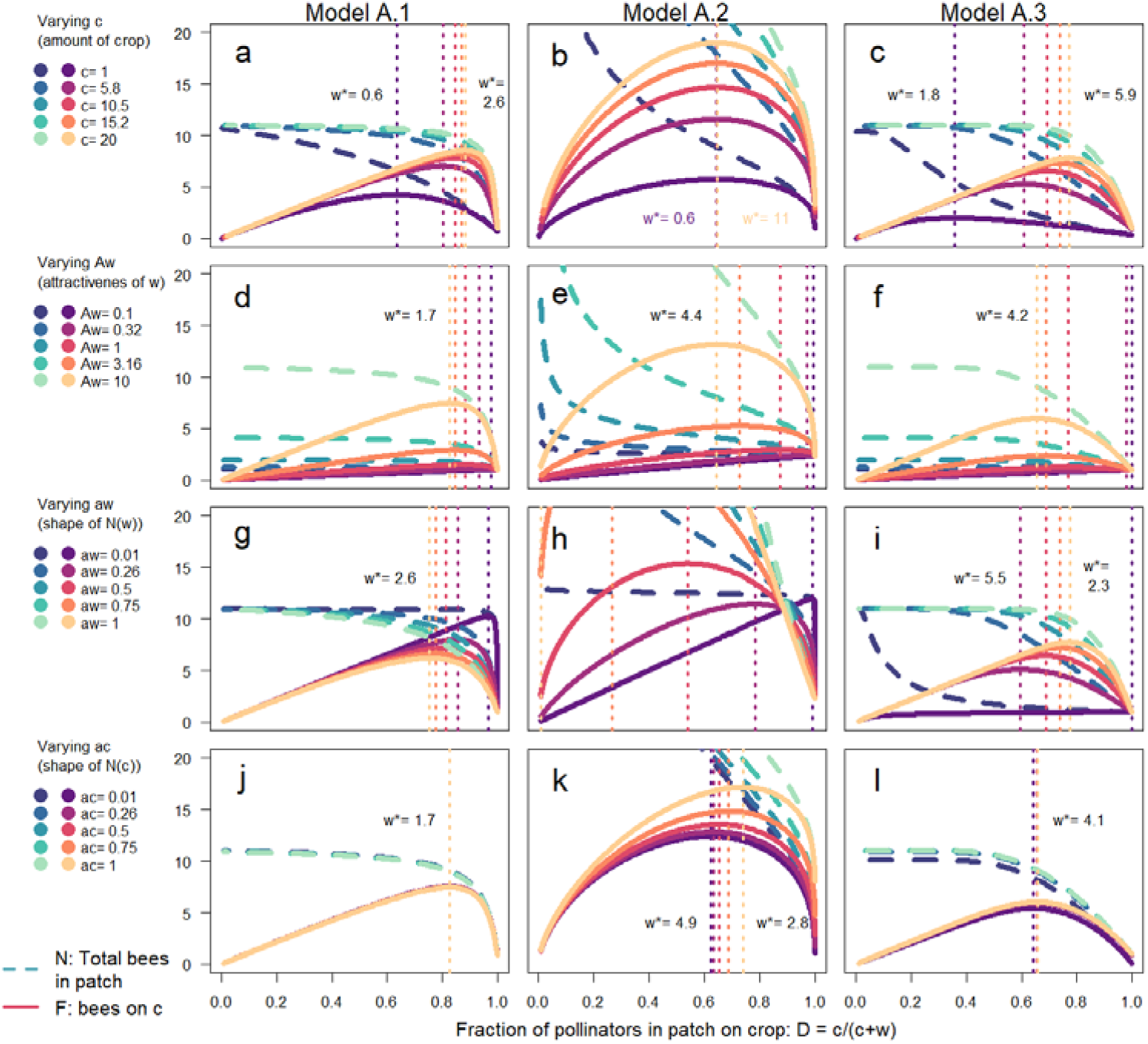
The behaviour of N (blue shaded, dashed lines) and F (red shaded, solid lines) across the three models with differing parameters (a, c, g: model A.1; b, d, h: model A.2; c, e, i: Model A.3). We show *D* on the x-axis, which is the empirically measurable fraction of pollinators in the patch that visit *c* instead of *w*; as *w* increases, *D* decreases. Default parameters used in this plot are *c*=8, *A*_w_=10, *A*_c_=1, *a*_w_=*a*_c_=0.4. Note that since *c* is fixed in panels d-l, each *D* corresponds to the same *w* across all parameter values in all panels but the first row. However, in panels a-c, as *c* is varied, the same *D* can correspond to different *w*. In model A.2 (panel b), *D*(*w\**) stays constant across different *c*, but this implies a changing *w\** along with *c* (labeled in color for max. and min. *c*). *D*(*w\**) is only invariant with *c* if *a*_w_=*a*_c_.

### 2. Model B: effects of parameters on attraction and choice for fixed area

Model B in the main manuscript (eqs. [10-13]) differs from the models A above in that some resource other than pollinators is assumed to be limiting (e.g. total area), and that thus an increase in *c* implies a decrease in *w* and vice versa. Using numerical calculations, we explored the effects of different parameter values on the optimal ‘amount’ of wildflowers, from the point of view of maximizing the number of pollinators on *c*. Because of the decelerating relationship of *N* with density of either species, the total number of bees in the patch is always highest at an intermediate amount of *c* and *w* (hump-shaped *N* lines in Fig. S3), as long as either *b*≥1 or *A*_w_≥1. As in the previous models, high attractiveness of *w* (attraction of pollinators to the patch), *A*_*w*_, generally has a large and direct positive effect on the total number of pollinators and thus also on *p\** (Fig. S3c and d). While *A*_*w*_ reflects how well wildflowers attract bees to the patch, *b* is the parameter that measures how much bees choose to visit wildflowers once in the patch, in relation to how much area the wildflowers occupy. If the limiting resource is not area, *b* still reflects the value to bees produced by *w* for the amount of the limiting resource consumed. If *b*=1, the value of *c* and *w* per area is the same (Fig. S3 panels c-h and in panels a, b where *b*=1). If b<1, wildflowers are not as valuable to bees as the crop flowers are, for the same area occupied; as a consequence, additional wildflowers do not increase pollinators present and do not facilitate crop pollination. The strongest facilitation for the crop is achieved with highly attractive wildflowers that have both a high *b* and *A*_w_. Perhaps counterintuitively, such flowers need not compete with the crop for pollinators but instead lead to substantial pollinator attraction generated for the area available, which can benefit the crop.

**Fig. S3:**
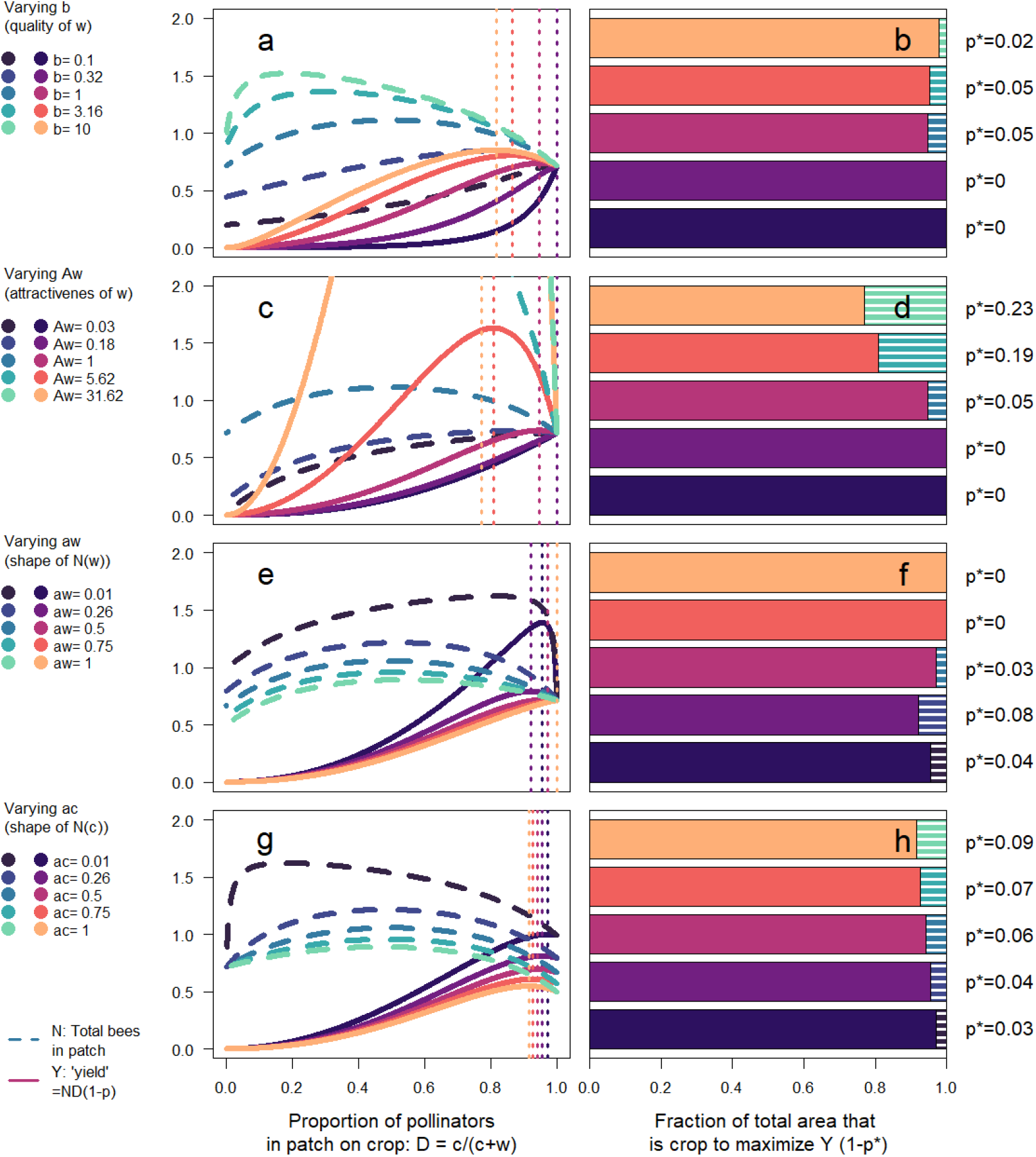
The behaviour of model B, where the total of *c* + *w* is limited by a shared resource (such as area). Panels a,c,e,g are analogous to fig. S2: blue shaded lines show *N*, the total number of pollinators in the patch; red shaded lines show yield on *c* at different parameter values. Panels b, d, f, h show the proportion of total area that is occupied by crop at maximal yield, i.e. 1-*p\** (red shaded bars; the rest of the total area is occupied by wildflowers, *p\**). Top row: varying *b*, the inherent value of the wildflowers to pollinators compared to that of the crop, i.e. affecting the choices of bees in the patch. Second row: varying *A*_w_, the attraction of pollinators to the patch by the wildflowers. Third and Fourth row: varying *a*_w_ and *a*_c_, respectively, parameters determining the shape of *N*(*c*,*w*)). Default parameter values are *b*=1, *A*_w_=*A*_c_=1, *a*_w_=*a*_c_=0.4.

